# Genome-based transmission modeling separates imported tuberculosis from recent transmission within an immigrant population

**DOI:** 10.1101/226662

**Authors:** Diepreye Ayabina, Janne O Rønning, Kristian Alfsnes, Nadia Debech, Ola B Brynildsrud, Trude Arnesen, Gunnstein Norheim, Anne-Torunn Mengshoel, Rikard Rykkvin, Ulf R Dahle, Caroline Colijn, Vegard Eldholm

**Author notes:** corresponding authors Vegard Eldholm - Caroline Colijn.

## Abstract

**Background:** In many countries tuberculosis incidence is low and largely shaped by immigrant populations from high-burden countries. This is the case in Norway, where more than 80 per cent of TB cases are found among immigrants from high-incidence countries. A variable latent period, low rates of evolution and structured social networks make separating import from within-border transmission a major conundrum to TB-control efforts in many low-incidence countries.

**Methods:** Clinical *Mycobacterium tuberculosis* isolates belonging to an unusually large genotype cluster associated with people born in the Horn of Africa, have been identified in Norway over the last two decades. We applied modeled transmission based on whole-genome sequence data to estimate infection times for individual patients. By contrasting these estimates with time of arrival in Norway, we estimate on a case-by-case basis whether patients were likely to have been infected before or after arrival.

**Results:** Independent import was responsible for the majority of cases, but we estimate that about a quarter of the patients had contracted TB in Norway.

**Conclusions:** This study illuminates the transmission dynamics within an immigrant community. Our approach is broadly applicable to many settings where TB control programs can benefit from understanding when and where patients acquired tuberculosis.

## Introduction

The ‘End TB strategy’ of the World Health Organization aims to reduce the global incidence of tuberculosis (TB) to 100 cases per million by 2035. Reaching this goal entails reducing the global TB incidence to the rates currently seen in countries with the lowest incidence, whereas low-incidence countries must aim for further reduction in incidence levels.

As the TB epidemic is fading out of the local population, the TB incidence in Norway now largely reflects the level of immigration from high-TB-incidence countries [3, 12, 5]. With effective case finding and case management, TB transmission from immigrant populations to the Norwegian-born population has been found to be very limited [4], but it does occasionally occur [11]. In low-incidence countries, preventing transmission within immigrant groups originating from high-incidence countries is of utmost importance if the overall TB incidence is to be reduced further. Detecting transmission within immigrant populations remains a complicated task as it requires the ability to distinguish between import and recent transmission in the receiving country. Even when genome sequences are available for analysis, the reconstruction of detailed transmission histories is far from straightforward. As a result of a low evolutionary rate and highly variable and stochastic latency periods, this is especially true for TB [11, 7]. This is further complicated by social networks that are not randomly formed, and among other things shaped by shared cultural and ethnic backgrounds [18]

Several approaches have been developed to make use of genomic data to infer infector/infectee relationships. They differ in their statistical approaches, the complexity of the underlying epidemiological models and the assumptions they make about unsampled cases, transmission bottlenecks, pathogen evolution, diversity inside hosts and the likelihood of transmission events. Key parameters that must be specified or estimated include the generation time (time from an individual becoming infected until infecting others) and patient and health system delay (time from symptom onset to diagnosis). In addition, inference requires timing information and a model of the pathogen’s evolution to connect the acquisition of polymorphisms to the (variable and uncertain) time that has elapsed. [8, 16, 19, 17, 6].

Didelot et al. developed a method to infer transmission networks from time-labelled phylogenies assuming a Susceptible-Infectious-Removed (SIR) epidemiological model [8]. Building on the same approach, Eldholm et al. implemented a latent state (“Exposed” category) in a similar framework to estimate the probability of pairs of patients being linked by a transmission event [11]. TransPhylo [7] is a Bayesian method for inference of transmission trees that also accounts for unsampled cases and infers transmission events and their timing given mutational events captured in a time-labelled phylogeny; it uses a branching model rather than an SIR or SEIR framework.

Here, we apply a novel approach to investigate whether immigrants diagnosed with TB represent imported cases or local transmission. We focused on a large genotypic *M. tuberculosis* cluster (Norwegian-African large Lineage 3 cluster; NAL3C), strongly associated with immigrants from the Horn of Africa, that has been identified in Norway consistently for almost 20 years [13]. Building on a temporal phylogeny built from genome-wide SNPs, we infer the posterior distribution of infection times *T_inf_* using TransPhylo. We then compare these with the time of arrival of individual patients in Norway, to ascertain probabilistically whether they became infected before or after their arrival in Norway. We confirm that most TB patients were indeed infected prior to arrival, but show that about 25% of the patients likely contracted TB after arrival in Norway.

## Results

Based on 24-loci Mycobacterial Interspered Repetitive Unit (MIRU) typing, 129 clinical *M. tuberculosis* NAL3C isolates from 127 patients, collected between 1997 and 2015 (Fig. 1A and 1B) were sequenced at the NRLM. A total of 1418 variable sites were identified, resulting in a mean pairwise-SNP distance of 43.22 separating the NAL3C isolates. A temporal phylogeny was estimated in BEAST 1.8.4 [9] utilizing sampling dates for temporal calibration (Fig. 1C).

**Figure 1:**
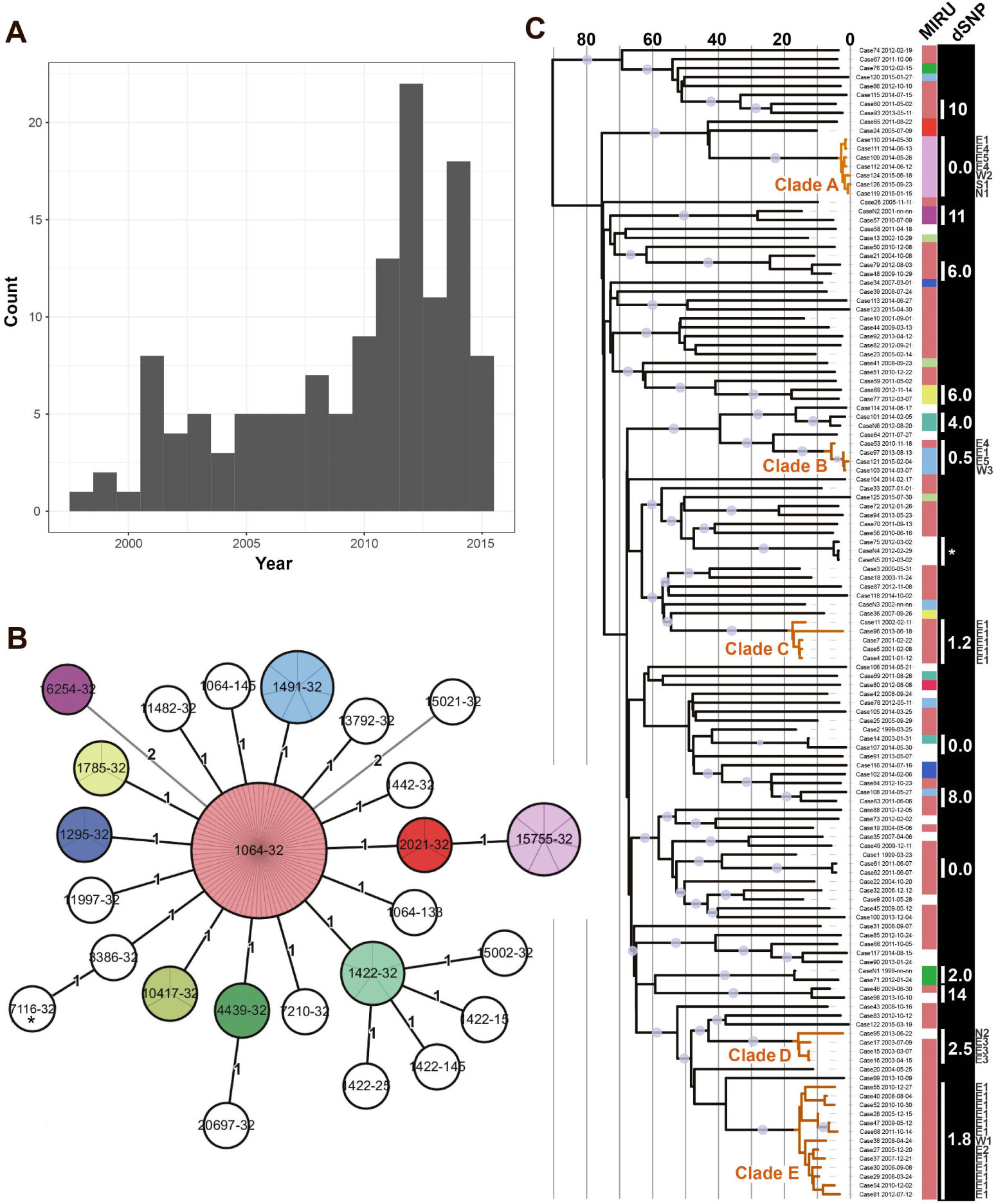
Clinical M. tuberculosis isolates and phylogenetic reconstruction. (A) Histogram illustrating sampling times for NAL3C isolates. (B) NAL3C minimum-spanning tree based on 24-loci MIRU genotypes. MIRU profiles identified in more than one isolate were assigned individual colors. (C) Temporal phylogeny constructed from genome-wide SNPs. The color strip next to the phylogeny indicates MIRU genotype, whereas the dSNP column denotes mean pairwise-SNP distances within each cluster. Clades amenable for TransPhylo transmission reconstruction are highlighted in orange. The time-axis on the phylogeny corresponds to years before 2015. An asterisk denotes three samples isolated from the same patient. Grey dots on branches indicate posterior probability of > 0.8. The county of residence of patients belonging to clades A-E is annotated in the right margin (see methods for details).

Comparing the MIRU-based minimum-spanning network and the whole-genome phylogeny of our samples, it was clear that the true genomic diversity of NAL3C was not at all captured by MIRU typing. Strikingly, we also find that the micro-evolution of MIRU loci within the cluster evolved in a way that was not informative for molecular epidemiological purposes. In fact, a number of homoplasic events led to the repeated evolution of identical MIRU types across the NAL3C (Fig. 1). Based on these analyses it is clear that MIRU typing worked rather well for crude grouping of isolates (*i.e*. to define inclusion criteria for whole genome sequencing), but that micro-evolutionary events, such as the mutation of a single MIRU locus, are not necessarily informative for molecular epidemiological inference. NAL3C belongs to lineage 3, an understudied *M. tuberculosis* lineage. Whether the mode and rate of MIRU evolution differs between lineages is a question that deserves attention, and indeed there is evidence that lineage strongly affects how well MIRU similarity predicts whole-genome sequence similarity [20]; this clearly affects the interpretation of MIRU data.

The high genetic diversity within the NAL3C cluster, combined with an overall phylogenetic structure characterized by multiple long terminal branches interspersed by a handful of tight clusters, suggested that the clinical TB cases in Norway represented samples drawn from a larger population of mainly unsampled cases presumably circulating in the Horn of Africa. Applying the inclusion criteria described in the Methods section resulted in a total of five clades (clades *A,B,C,D* and *E* shown in Figure 1) meeting the criteria for detailed transmission modelling. Most of the cases in these clades come from countries in the horn of Africa, two from Sudan and one case each from Ghana, Gambia, Iran, Thailand and Norway.

Figure 2 shows the arrival times of all cases from these clades for whom arrival times were retrievable (all in clades *A, B* and *E*), plotted on top of the posterior infection time distribution for each individual (see Methods and Supplementary Methods). It is quite clear that some of these patients (cases 30, 37, 40, 47, 54, 68 and 126) arrived in Norway before the estimated infection time (*T_inf_*). For all other cases, the estimated range for the time of infection has at least some overlap with the time of arrival. Using the posterior densities of infection times alongside the arrival times of the cases, we obtain probabilities of infection after the time of arrival in Norway (*P*(*t_arrv_*); see Table S3). The cumulative frequency plot of these probabilities (Fig. 3) shows that there are 16 cases with *P*(*t_inf_* after *t_arrv_*) > 0.5 and 12 cases with *P*(*t_inf_* after *t_arrv_*) > 0.9.

**Figure 2:**
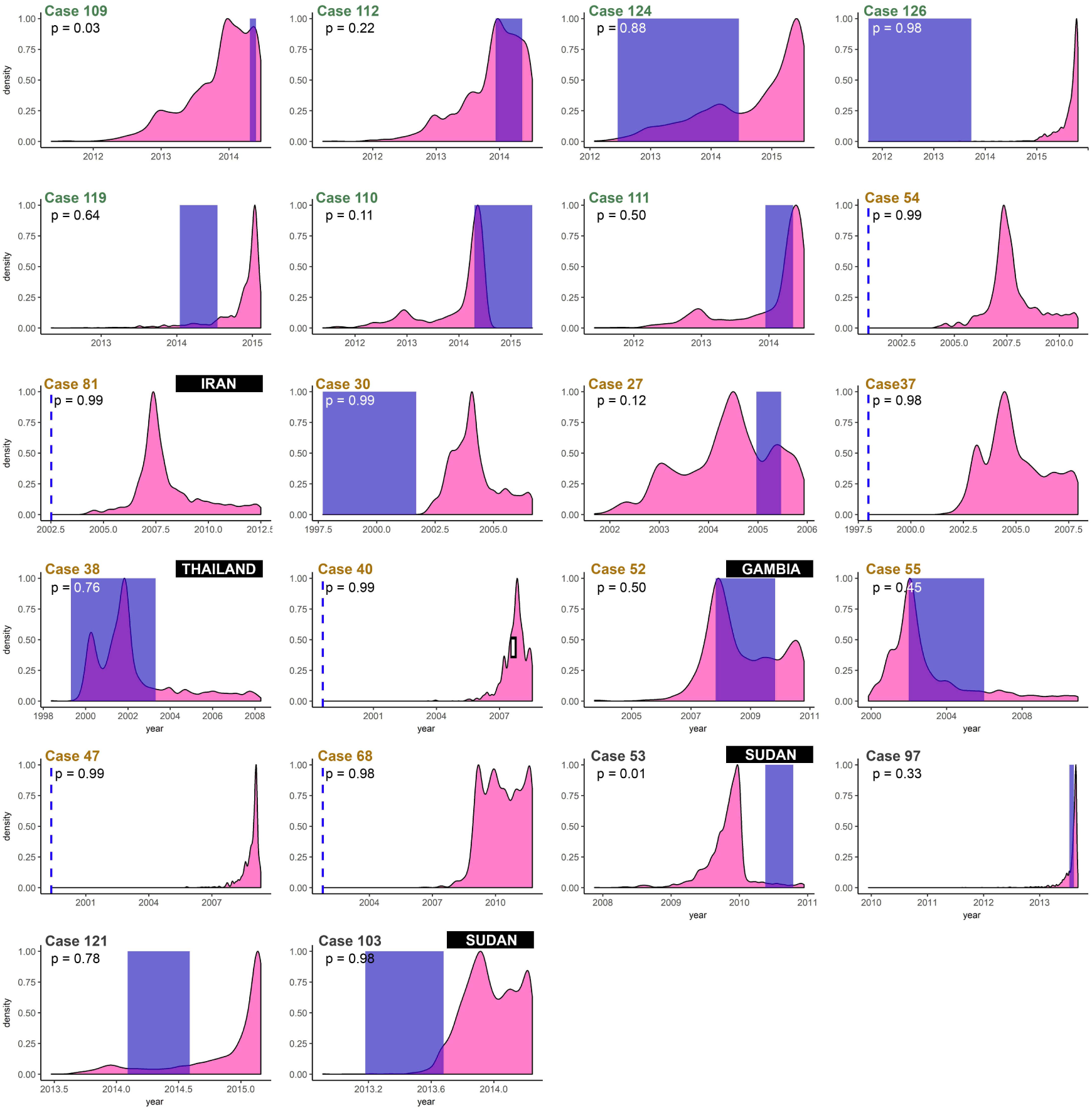
Arrival times (in blue) plotted with estimated infection times for all cases of interest with available data. The case numbers are colored by clade assignment (clade A in green, clade E in orange and clade B in grey). The blue shaded area covers the time from earliest and latest possible arrival times, whereas a dotted single line indicates the latest possible arrival time. The listed probabilities P indicate the probability of infection after arrival in Norway, averaged over 10 different TransPhylo inference procedures. The country of origin of patients not originating from the Horn of Africa is annotated in black boxes.

**Figure 3:**
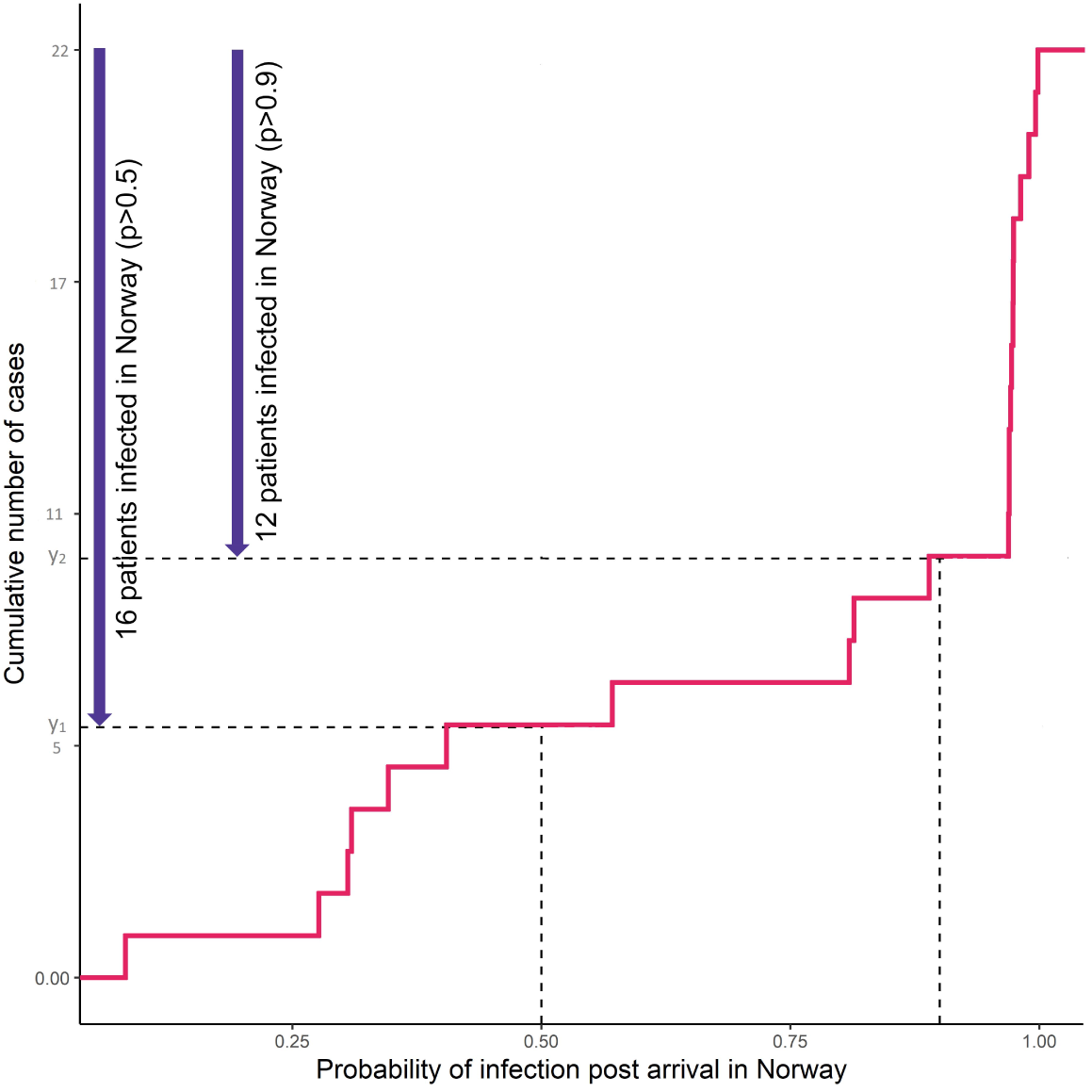
Probability of infection after arrival in Norway for 22 cases included in TransPhylo analyses. The lines annotate the number of cases with probabilities equal to 0.5 and 0.9 in the cumulative distribution plot.

In clades *A, B* and *E*, cases 28 and 29 lacked arrival time information. For case 29 (Somali) we could conclude that the patient likely contracted TB in Norway, as the inferred infector was also infected in Norway. The genome of the isolate from Case 28, an immigrant from Ghana, was identical to the one from case 29, indicating that case 28 was also infected in Norway. Next, we looked into clades *C* and *D* for which arrival information was lacking for all isolates. For clade *C*, TransPhylo inferred that the same unsampled case had infected both a Norwegian, Ethiopian and two Somali patients, as well as a final Somali patient via another unsampled intermediate (see Figure S7), though we note that with long and variable infectious periods, as is the case for TB, considerable uncertainty remains in the details of reconstructed transmission events. However, our results suggest that all five patients contracted TB in Norway. In clade *D*, all patients were Somali. This, combined with a lack of arrival information for these patients, makes it impossible to distinguish between transmission before or after arrival.

Finally, in order to obtain a more complete picture of transmission in Norway, beyond clades that were amenable to transmission inference using TransPhylo, we manually investigated the temporal phylogeny for pairs and triplets of closely related isolates for evidence of transmission in Norway. Following the inclusion criteria applied for TransPhylo inference, we only included pairs and triplets with an estimated most recent common ancestor after 1995. Based on a combination of arrival times, disease manifestation and country of origin, we were able to identify another three instances of very probable transmission in Norway. For five cases we could conclude that transmission in Norway was highly unlikely, whereas no conclusion could be drawn for six of the cases (see Tables S4 and S5 for a summary of the evidence).

In total, we conclude that 26 out of 129 NAL3C cases were probably infected in Norway. For 10 cases we were unable to conclude, whereas the remaining 91 probably represented instances of imported TB (see Table 1).

**Table 1:** Summary of transmission inference.

| Inference type | Yes | Undetermined | No | Clades |
| --- | --- | --- | --- | --- |
| TP (with arrival info) | 16 | 0 | 6 | A,B,C |
| TP (without arrival info) | 7 | 4 | 0 | C,D,E |
| Smaller clusters | 3 | 6 | 5 | – |
| Other cases | 0 | 0 | 80 | - |
| Total (%) | 26(20) | 10(8) | 91(72) | - |

TP = TransPhylo inference

Retrospectively, we retrieved available contact tracing information and extended epidemiological data for cases belonging to clades *A, B* and *E*. The data were incomplete, but useful information was available for six of the cases. For five of these patients the extended data supported our inference: Three cases estimated as likely infected in Norway had known TB contacts in Norway (cases 40, 47 and 81); The estimated transmission time concurred with an earlier episode of tuberculosis (case 55); or a negative TB screen upon arrival in Norway was consistent with our inference of infection after arrival (case 103). However, in once case (case 119) we estimated infection in Norway but the patient reported to have had an episode of symptomatic, untreated TB before arrival; the recent isolate could reflect a re-infection, or our estimation could be incorrect.

## Discussion

There are a number of limitations to our approach. While we took steps to accommodate phylogenetic uncertainty by repeating our analysis starting with different input trees, we do not have a way to compare parameters like *N_e_g* directly to data, nor can we directly validate the inferred times between infection and sampling, or between infection and infecting others, against data. This is a fundamental limitation in any outbreak reconstruction task, as the ground truth is not known. However, our results on the timing of infection are robust to different input phylogenies and choices of priors. Similarly, we do not know the contribution of smear-negative individuals to transmission, nor, indeed, the constancy of smear status over the course of infection. We used a penalty on transmission by smear-negative cases because they are widely believed to be less infectious than smear-positive individuals; our conclusions that there is evidence of TB transmission in Norway, and for whom, are not affected by removing this penalty.

We do not have individual-level data such as factors that may have promoted infection in a short time frame, so we cannot draw conclusions about host factors that may have caused rapid progression to disease. We also note that by the nature of the study, we are not finding and including individuals who have “latent” (asymptomatic, contained) tuberculosis; only those with active disease would produce culture-positive samples that can be sequenced, and TransPhylo only augments the tree with unsampled individuals if these are in transmission chains at the current MCMC state. Accordingly, the generation times in the model are reflective of generation times for individuals who progress to active disease by the time of sampling, and who have done so in time to infect others who have progressed to active disease and arisen in transmission chains in our inference. Finally, TransPhylo does not model a contraction of generation times as an epidemic proceeds [15]; the branching process model is built on the approximation that clustering is low (individuals are unlikely to expose each others’ contacts) and that the susceptible population size is much larger than the infected population. Compared to currently available survival analysis methods that do not require these assumptions, TransPhylo carries the advantages that it handles both the possibility of unsampled individuals and in-host diversity.

Even without explicitly reconstructing transmission events, some conclusions about the timing of transmission can be drawn from the timed phylogenetic tree. Each tip of the tree corresponds to a different host, which implies that there must be at least one transmission event on the path between every pair of tips [8, 7]. Recent branching in the timed phylogeny therefore indicates recent infection. Our analysis goes further, by producing probability distributions for infection times for individual patients; this was achieved by combining genome sequences, clinical data and epidemiological data to inform transmission modeling. We believe this work represents a conceptually novel and useful approach to tackle an important public health issue in low TB incidence countries.

Five of the patients belonging to potential transmission clusters and for whom arrival times were known originated from countries other than those in the Horn of Africa (Fig. 2). Four of these, originating from Thailand, Iran, Sudan and the Gambia, were inferred to have been infected in Norway (*p* ≥ 0.5). The fifth patient was from Sudan, and was inferred to have been infected prior to arrival. The *M. tuberculosis* isolates from two additional Sudanese patients (Case 24 and 83) were found alone on long terminal branches of the phylogeny (Fig. 1), also suggesting that they were independently imported cases. Two Tanzanian patients also represented relatively clear import events with clinical isolates situated on long terminal branches (Case 33 and Case 57). Apart from patients from the Horn of Africa, the only foreign patients we could conclude had been infected prior to arrival originated from Tanzania and Sudan. Sudan shares a border with Eritrea and Ethiopia whereas Tanzania lies to the south of Somalia and Ethiopia, with Kenya in between. Taken together, these findings suggest that the geographic range of the NAL3 cluster stretches beyond the Horn of Africa into neighboring countries including Sudan, Kenya and Tanzania.

We analyzed our inferences in the context of where individual patients lived in Norway at the time of diagnosis. The seven patients in Clade A were all of Eritrean origin (table S3), but were scattered between six different counties covering large parts of Norway at the time of diagnosis. This suggests that those patients inferred to have been infected in Norway were likely infected en route or soon after arrival to a common destination in Norway such as a reception center. Unfortunately no information is available to clarify this. The four patients in Clade B were from three nations (Table S3) and lived in four different counties, again suggesting a reception center as a likely place of infection. Nine out of 11 patients in Clade E lived in the same county (E1) and one in a neighboring county (E2), suggesting transmission in the community. The five patients in clade C were from three different nations, and all lived in county E1 at the time of diagnosis, supporting our inference of transmission in Norway (Table S5). Finally, in clade D, where no conclusions regarding transmission in Norway could be made (Table S5), three of four patients lived in county E3, which does suggest that transmission in Norway could be responsible for some of the cases.

We find that 26 patients belonging to the NAL3C cluster were most probably infected in Norway, whereas 91 patients had probably contracted TB prior to arrival in Norway. For 10 patients, our analyses were inconclusive. In addition, we were unable to retrieve samples for DNA extraction for three patients, originating from Norway, the former Yugoslavia and Namibia respectively. Based on their country of origin alone, these were all probably infected in Norway. Only considering patients for which a conclusion could be drawn, about a quarter were most likely to have contracted TB in Norway, a far from trivial proportion. It should be noted that some patients might also have been infected during travels to the country of origin after arrival in Norway, but we were not in a position to investigate this possible order of events. A very recent study from the Netherlands and Denmark identified the 1064-32 MIRU-type among refugees and immigrants with a similar country-profile as observed in Norway [14]. Whole genome sequencing of 40 isolates revealed a pairwise SNP-distance of 80, almost twice as high as observed here. Although formal transmission reconstruction was not performed, the high diversity supports their conclusion that transmission in the Netherlands and Denmark is very limited. The lower diversity in Norway also suggests that recent transmission in the country has affected the observed population structure of NAL3C.

In Norway, contact tracing is initiated around all pulmonary TB cases, but the intensity of the effort is higher when recent transmission is suspected. The bar for initiating broad contact tracing efforts will thus often be higher for TB-cases belonging to high-risk groups such as immigrants from high-incidence countries. However, we found that there is substantial evidence of ongoing transmission in Norway. Our findings thus highlight the importance of active contact-tracing in high-risk groups in low-incidence countries.

## Conclusion

Transmission modeling based on a combination of clinical, epidemiological and genomic data, can assist public health authorities in understanding where and when patients are infected, and can aid in the design of appropriate TB control measures. Importantly, the use of genomic and epidemiological to quantify local transmission can thus serve as a useful metric to gauge the quality of local/national TB programmes.

## Methods

### Sample collection and epidemiological data

The NRLM maintains a national culture collection consisting of all culture-positive TB cases in Norway and is responsible for susceptibility testing and genotyping. From 1997 till 2010 *IS6110* restriction fragment length polymorphism (RFLP) was the routine method for molecular epidemiological surveillance. In this period, a large *IS6110*-RFLP-cluster associated with patients from the Horn of Africa was identified [13]. Following the replacement of *IS6110*-RFLP typing with 24-loci MIRU-genotyping in 2011, these isolates were re-typed and the majority of isolates belonged to the MIRU type 1064-32 following on the MTBC 15-9 nomenclature [2]. All subsequent *M. tuberculosis* isolates have been MIRU typed. In order to study the transmission dynamic of this cluster, we included all isolates sampled between 1997 and 2015 that differed at zero to two loci relative to the 1064-32 genotype. In total, 133 isolates matched the inclusion criteria, of which 130 could be retrieved and were submitted to whole-genome sequencing on the Illumina platform. Initial analyses revealed one of these to be a clear outlier only distantly related to the other 129 isolates; it was thus excluded. The 129 isolates represented 127 patients (three isolates were from the same patient). The cluster, as defined by *IS6110*-RFLP genotyping, was recently termed “Cluster X” [13]. However, we coined the more informative term Norwegian-African large lineage 3 cluster (NAL3C) for the current study. Minimal epidemiological data were extracted from the Norwegian Surveillance System for Communicable Diseases (MSIS), which stores clinical and epidemiological data on all TB cases notified by clinicians. The study protocol was approved by the Regional Ethics Committee (reference 2015/2127).

### Variant calling and phylogenetic analyses

DNA was extracted from *M. tuberculosis* grown on Lowenstein-Jensen slants as described previously [10]. Paired-end sequences were generated on the Illumina MiSeq and NextSeq platforms (250 and 150 bp read length respectively). High quality single nucleotide polymorphisms were identified following the same procedures as described in [10]. After removal of the single outlier isolate, this resulted in 1418 variable sites that were used for evolutionary analyses as outlined below. Median sequencing depth of the 129 genomes ranged from 20x to 161x. All sequence reads are available under ENA study accession PRJEB23495. Individual run accessions, sampling years and MIRU data for all NAL3C isolates are listed in dataset S01. Marginal likelihood estimates in Beast 1.8.4 [9] were performed to identify the optimal substitution, clock and demographic models for Bayesian evolutionary analyses. A GTR model with relaxed clock combined with a Skyride demographic model was favored.

### Transmission reconstruction

The maximum credibility tree of the 129 isolates is characterized by long branch lengths with a few clades that have relatively short branch lengths (Fig. 1). As the branch lengths of a timed phylogenetic tree represent duration of evolution [1], we assumed that these clades represent densely sampled clusters of cases whereas the long branches represent cases with unsampled infectors. We thus reasoned that putative transmission clusters in Norway would be represented by sub-clades of closely related isolates within the larger NAL3C cluster. As the vast majority of immigrants from the Horn of Africa came to Norway after 1995, following the withdrawal of UN from Somalia, we only considered clades with an inferred most recent ancestor younger than 20 years (corresponding to 1995). As a tree must include at least four cases for meaningful modeling in TransPhylo, this inclusion criterion was also applied.

A total of five clades (clades *A,B,C,D* and *E* shown in Figure 1, met the criteria for detailed transmission modeling. There are a total of 33 cases in the selected clades, with times of arrival in Norway available for 22 of them. In addition, manual transmission inference was performed on pairs and triplets of closely related isolates. TransPhylo requires priors for the generation time and time to sampling; these were estimated from data using all cases in one of the clades (Clade A); the fitted posterior values were used to specify the generation time and sampling time densities in the onward analysis. We explored a range of prior distributions (Table S2). Within this range the key results about recent transmission are not dependent on the choice of prior. To reflect that fact that smear-negative patients are likely to be less infectious than smear-positive patients, we modelled transmissions from smear-negative patients as 75% as likely as transmissions from smear-positive patients (all else being equal); results on recent transmission are not altered when this penalty is removed.

TransPhylo is a “two-step” method, relying on first constructing a timed phylogenetic tree and then performing Bayesian inference of the transmission events. We accounted for phylogenetic uncertainy by repeating the inference on different posterior timed phylogenetic trees (see Supplement). TransPhylo requires assumptions about the parameters *r* and *p* (negative binomial offspring distribution parameters), the in-host effective population size *N_e_g* and the overall sampling probability *pi*. We fixed *N_e_g* assumed a mean offspring of 1 and a sampling fraction of 0.95 (See Supplement).

### Geographic information in Norway

Norway is divided into 19 counties. Here we do not use county names in order to minimize the use of possibly sensitive data, but rather refer to these counties as E (East), S (South), W (West) and N (North) followed by a number unique to each county.

Please see the Supplementary Methods for a full description of all methods employed.

## Acknowledgements

We would like to acknowledge the Norwegian Sequencing Center for sequencing a subset of the samples on the NextSeq platform and the staff at the National Reference Laboratory for Mycobacteria, Norway, for maintenance and genotyping of a national TB culture collection. This work was supported by the Engineering and Physical Sciences Research Council of the United Kingdom (EPSRC EP/N014529/1 and EPSRC EP/K026003/1 - Caroline Colijn) and the Nigerian Universities Commission (under the presidential special scholarship scheme for innovation and development (PRESSID - Dipreye Ayabina))

## References

[1] D. Alexei and B. Remco. Bayesian evolutionary analysis with BEAST. Cambridge University Press, 2015.

[2] C. Allix-Béguec, M. Fauville-Dufaux, and P. Supply. Three-year population-based evaluation of standardized mycobacterial interspersed repetitive-unit-variable-number tandem-repeat typing of Mycobacterium tuberculosis. Journal of Clinical Microbiology, 46(4):1398–406, Apr 2008.

[3] M. Carballo and M. Mboup. International migration and health: A paper prepared for the policy analysis and research programme of the global commission on international migration. 2005.

[4] U. R. Dahle, V. Eldholm, B. A. Winje, T. Mannsåker, and E. Heldal. Impact of immigration on the molecular epidemiology of *Mycobacterium tuberculosis* in a low-incidence country. American Journal of Respiratory and Critical Care Medicine, 176(9):930–935, nov 2007.

[5] U. R. Dahle, P. Sandven, E. Heldal, and D. A. Caugant. Continued low rates of transmission of *Mycobacterium tuberculosis* in Norway. Journal of Clinical Microbiology, 41(7):2968–73, Jul 2003.

[6] N. De Maio, C.-H. Wu, D. J. Wilson, S. Gaudieri, S. Pham, and A. Chopra. SCOTTI: Efficient reconstruction of transmission within outbreaks with the structured coalescent. PLOS Computational Biology, 12(9):e1005130, Sep 2016.

[7] X. Didelot, C. Fraser, J. Gardy, and C. Colijn. Genomic infectious disease epidemiology in partially sampled and ongoing outbreaks. Molecular Biology and Evolution, 195(4):msw075, Jan 2017.

[8] X. Didelot, J. Gardy, and C. Colijn. Bayesian inference of infectious disease transmission from whole-genome sequence data. Molecular Biology and Evolution, 31(7):1869–79, Jul 2014.

[9] A. J. Drummond, M. A. Suchard, D. Xie, and A. Rambaut. Bayesian Phylogenetics with BEAUti and the BEAST 1.7. Molecular Biology and Evolution, 29(8):1969–1973, Aug 2012.

[10] V. Eldholm, J. H.-O. Pettersson, O. B. Brynildsrud, A. Kitchen, E. M. Rasmussen, T. Lillebaek, J. O. Rønning, V. Crudu, A. T. Mengshoel, N. Debech, K. Alfsnes, J. Bohlin, C. S. Pepperell, and F. Balloux. Armed conflict and population displacement as drivers of the evolution and dispersal of *Mycobacterium tuberculosis*. Proceedings of the National Academy of Sciences of the United States of America, 113(48):13881–13886, Nov 2016.

[11] V. Eldholm, A. Rieux, J. Monteserin, J. M. Lopez, D. Palmero, B. Lopez, V. Ritacco, X. Didelot, and F. Balloux. Impact of HIV co-infection on the evolution and transmission of multidrug-resistant tuberculosis. eLife, 5:e16644, Aug 2016.

[12] M. Farah, H. Meyer, R. Selmer, E. Heldal, and G. Bjune. Long-term risk of tuberculosis among immigrants in Norway. International Journal of Epidemiology, 34(5):1005–1011, Jul 2005.

[13] B. R. Guzman Herrador, K. Rønning, K. Borgen, T. Mannsåker, and U. R. Dahle. Description of the largest cluster of tuberculosis notified in Norway 1997–2011: is the Norwegian tuberculosis control programme serving its purpose for high risk groups? BMC Public Health, 15(1):367, Dec 2015.

[14] R. Jajou, A. de Neeling, E. M. Rasmussen, A. Norman, A. Mulder, R. van Hunen, G. de Vries, W. Haddad, R. Anthony, T. Lillebaek, W. van der Hoek, and D. van Soolingen. A predominant VNTR cluster of Mycobacterium tuberculosis isolates among asylum seekers in the Netherlands and Denmark deciphered by whole genome sequencing. Journal of Clinical Microbiology, Nov. 2017.

[15] E. Kenah, T. Britton, M. E. Halloran, I. M. Longini, T. Leitner, and V. de Jager. Molecular Infectious Disease Epidemiology: Survival Analysis and Algorithms Linking Phylogenies to Transmission Trees. PLOS Computational Biology, 12(4):e1004869, Apr 2016.

[16] M. Kendall, D. Ayabina, and C. Colijn. Estimating transmission from genetic and epidemiological data: a metric to compare transmission trees. Sep 2016.

[17] D. Klinkenberg, J. A. Backer, X. Didelot, C. Colijn, J. Wallinga, and D. Haydon. Simultaneous inference of phylogenetic and transmission trees in infectious disease outbreaks. PLOS Computational Biology, 13(5):e1005495, May 2017.

[18] A. Sandgren, M. S. Schepisi, G. Sotgiu, E. Huitric, G. B. Migliori, D. Manissero, M. J. van der Werf, and E. Girardi. Tuberculosis transmission between foreign- and native-born populations in the EU/EEA: a systematic review. The European Respiratory Journal, 43(4):1159–71, Apr 2014.

[19] C. J. Worby, M. Lipsitch, and W. P. Hanage. Shared genomic variants: identification of transmission routes using pathogen deep sequence data. American Journal of Epidemiology, Jun 2017.

[20] D. Wyllie, J. Davidson, T. Walker, P. Rathod, T. Peto, E. Robinson, D. W. Crook, G. Smith, and C. Campbell. A quantitative evaluation of MIRU-VNTR typing against whole-genome sequencing for identifying Mycobacterium tuberculosis transmission: A prospective observational cohort study. Jan. 2018.

